# Environmental gradients mediate dispersal evolution during biological invasions

**DOI:** 10.1101/2023.12.08.570855

**Authors:** John W. Benning, Eliza I. Clark, Ruth A. Hufbauer, Christopher Weiss-Lehman

## Abstract

Rapid dispersal evolution at the edge of a range expansion can accelerate invasions. However, expanding populations will often encounter environmental gradients that entail a fitness cost of dispersal. We used an eco-evolutionary model to explore how environmental heterogeneity influences adaptation and dispersal evolution during range expansion and, in turn, modulates the speed and predictability of invasion. Environmental gradients opposed the evolution of increased dispersal during invasion, even leading to the evolution of reduced dispersal along steep gradients. Counterintuitively, reduced dispersal allowed for faster expansion by minimizing maladaptation. While evolution across homogenous landscapes caused invasions to be highly unpredictable, even shallow environmental gradients greatly increased invasion predictability. We illustrate our model’s potential application to prediction and management of invasions by parameterizing it with data from a recent invertebrate range expansion. Overall, we find that environmental heterogeneity and local adaptation strongly modulate the effect of dispersal evolution on invasion trajectories.

## Introduction

The history of biodiversity on Earth is largely a story of invasion. From plants’ colonization of land 500 million years ago to contemporary range shifts with climate change, species’ geographic range expansions have repeatedly rearranged global biogeography. Though the timescale and details of these invasions vary greatly, they are underlain by the same fundamental process — the expansion of populations into previously unoccupied habitat. In recent decades, much attention has focused on the spread of introduced species as it became clear that anthropogenic movement of organisms could cause massive ecological and economic damage. Work has documented the ecological effects of invasive exotic species (Vilà *et al*. 2011), characterized traits that may promote invasiveness (Van Kleunen *et al*. 2010), and examined evolutionary processes during invasion (Urbanski *et al*. 2012; Colautti & Barrett 2013). Yet, predicting the spatio-temporal dynamics of invasion itself — namely, the rate of expansion — remains difficult (Miller *et al*. 2020). From a purely ecological perspective, expansion rate can be estimated quite simply based on two population parameters: intrinsic growth rate and dispersal ability (Hastings *et al*. 2004). However, empirical and theoretical work increasingly show that populations can evolve rapidly during invasion, which makes predicting invasion much more difficult (Burton *et al*. 2010; Colautti & Lau 2015; Fronhofer & Altermatt 2015; Phillips 2015; Williams *et al*. 2016; Ochocki & Miller 2017; Weiss-Lehman *et al*. 2017; Andrade-Restrepo *et al*. 2019; Peischl & Gilbert 2020; Urquhart & Williams 2021). Here we focus on two evolutionary processes that can modulate invasion dynamics on ecological timescales: dispersal evolution and local adaptation.

The evolution of heightened dispersal in leading edge populations during range expansion has been demonstrated in a variety of taxa in both experimental laboratory expansions and natural populations (Cwynar & MacDonald 1987; Simmons & Thomas 2004; Phillips *et al*. 2006; Hughes *et al*. 2007; Williams *et al*. 2016; Ochocki & Miller 2017; Weiss-Lehman *et al*. 2017; Van Petegem *et al*. 2018). This phenomenon is thought to be largely due to the process of spatial sorting, which arises from the simple fact that the most dispersive individuals will tend to aggregate at the invasion front. As long as dispersal traits are heritable, reproduction of these high-dispersal individuals will serve to increase dispersal ability at the invasion front compared to the range core (Shine *et al*. 2011). Spatial assortment of dispersal ability can be further augmented if individual fitness is negatively influenced by density (via intraspecific competition), such that low competition environments at the invasion front will result in high fitness of invaders. In this case, dispersal beyond the current range edge will be favored by natural selection due to an escape from competition (Travis *et al*. 2009; Perkins *et al*. 2013; Miller *et al*. 2020). Such increases in dispersal will increase invasion speed through time (Travis & Dytham 2002; Phillips *et al*. 2010; Perkins *et al*. 2013; Miller *et al*. 2020). But evolutionary processes acting on expanding populations do not only increase invasion speed — they are also expected to *decrease* the predictability of invasion. This is because evolution at the invasion front is likely to be highly influenced by stochastic processes due to increased genetic drift in small invading populations, which increases the variability in invasion rate across instances (Phillips 2015; Weiss-Lehman *et al*. 2017; Williams *et al*. 2019).

A second fundamental evolutionary process relevant to range expansions is adaptation. Given spatial environmental heterogeneity, natural selection will favor genotypes adapted to their local environment, leading to the widespread (though not ubiquitous) phenomenon of population local adaptation (Hereford 2009; Colautti & Barrett 2013; Briscoe Runquist *et al*. 2020; Gorton *et al*. 2022; Ittonen *et al*. 2022). In the context of range expansions, populations expanding across a landscape will often encounter environmental gradients — e.g., aridity, photoperiod, temperature — that result in phenotypic optima varying along the gradient. For instance, optimal phenology of bud break or flowering in plants often shifts across climatic gradients (Griffith & Watson 2006; Alberto *et al*. 2011; Gorton *et al*. 2022). Such gradients and the resulting spatial variation in selection will require continual adaptation for populations to spread across the gradient (Mayr 1963). Such local adaptation is found at similar rates within both native and exotic plant species (Oduor *et al*. 2016), indicating that spatially varying selection may influence the dynamics of many invasive species. As gradients steepen (i.e., phenotypic optima change more quickly across the landscape), both the fitness cost of dispersal and the “adaptive leap” required to colonize adjacent patches increases. Evolutionary models of range dynamics show that the steepness of such gradients is the key factor determining whether or not range expansion is possible for a species (Kirkpatrick & Barton 1997; Polechová & Barton 2015; Gilbert *et al*. 2017; Polechová 2018; Bridle *et al*. 2019).

While both dispersal evolution and local adaptation are expected to be common, we know little of how these two processes interact during range expansions. Intuition and theory suggest that, generally, they will oppose each other. Higher dispersal leads to increased gene flow between populations, which can stymie local adaptation and depress mean fitness when populations occupy environments with different selection regimes (Slatkin 1987; Lenormand 2002). Indeed, the idea of gene flow swamping adaptation underlies most evolutionary models of range limits (Haldane 1956; Kirkpatrick & Barton 1997; but see Polechová 2018). If there is spatially variable selection due to environmental heterogeneity, the negative effect of gene flow is thought to often lead to selection against dispersal as populations adapt to the local environment and the fitness cost of dispersal increases (Holt 1985; Billiard & Lenormand 2005; Ronce 2007; Bonte *et al*. 2012). Given these results, we might expect that selection along environmental gradients would constrain the evolution of heightened dispersal during invasion. Selection along gradients may also increase the predictability of invasion. Heterogeneity in habitat quality is expected to select against dispersal (Holt 1985), and thus environmental gradients might constrain increases in dispersal ability (and thus, expansion speed) that would be expected from spatial sorting. Furthermore, local maladaptation along a gradient is expected to slow range expansion, allowing more time for migrants to reach marginal populations, increasing population sizes at range edges (Gilbert *et al*. 2017). Such local selection could thus weaken the relative influence of genetic drift on evolution at the invasion front, potentially increasing the predictability of invasion (Gilbert *et al*. 2017; Williams *et al*. 2019; Miller *et al*. 2020).

Though scant, there are some empirical observations consistent with the idea that adaptation to novel environments can constrain the evolution of dispersal during range expansions. There was a reduced signal of dispersal evolution in flour beetle expansions across novel versus benign mesocosm landscapes (Szűcs *et al*. 2017; Weiss-Lehman *et al*. 2017), and dispersal evolution in a beetle species expanding south in the Western United states may be constrained by local adaptation to photoperiod (Clark *et al*. 2022, 2023). Theory has only recently begun to probe the interaction between dispersal evolution and local adaptation to gradients during range expansions and shifts. Andrade-Restrepo et al. (2019) showed that evolving dispersal induces changes in invasion tempo (wave vs. pulse) for a species expanding across an environmental gradient. Models of range shifts with climate change have shown that dispersal evolution can help rescue populations lagging behind shifting climatic isotherms (Weiss-Lehman & Shaw 2020; Block & Levine 2021). However, we do not know 1) how environmental heterogeneity regulates dispersal evolution during range expansion, or 2) how the process of local adaptation mediates the rate and predictability of invasion when dispersal can evolve.

We also have a limited understanding of how these spatial patterns of dispersal ability vary temporally — is elevated dispersal in invasion front populations permanent, or does the spatial pattern of dispersal vary through time? Travis and Dytham (2002) showed that when there was a cost to dispersal (modeled as a constant probability of dying when dispersing), heightened dispersal still evolved in sites at the front of the invasion wave, but once the wave passed, mean dispersal within a site decreased as the cost of dispersal became greater than the benefit of dispersal. Simmons and Thomas (2004) showed empirical evidence supporting this pattern of transient dispersal evolution in range expansions of bush crickets. However, if a dispersal cost arises indirectly due to maladaptation rather than as a direct, constant dispersal cost, it is unclear how we should expect the spatial patterns of dispersal evolution to change through time.

Overall, we lack a general understanding of how environmental gradients influence expectations for dispersal evolution and adaptation at the invasion front. Because of the potentially large effects of evolution on invasion rate and predictability, and the ubiquity of environmental gradients in nature, understanding the interplay of these forces would go far in building more predictive models of invasion dynamics. To that end, here we investigate the influence of an environmental gradient on the magnitude and temporal dynamics of dispersal evolution during invasion. Our individual-based simulation model allows for the evolution of two quantitative traits: a trait conferring adaptation to an environmental gradient, and a trait controlling dispersal ability. By varying the steepness of the environmental gradient and following the demography and evolution of populations expanding across it, we ask:

1. How does the severity of the environmental gradient modify dispersal evolution during expansion?
2. Does the relationship between dispersal evolution and predictability of range expansions depend on the severity of environmental gradients?
3. How does the interplay of dispersal evolution and local adaptation influence how long dispersal remains elevated after the invasion wave has passed?

## Material and methods

### Dispersal evolution and local adaptation during invasion

We used SLiM (Haller & Messer 2022), a forward-in-time population genetic modeling framework, to build a genomically-explicit, individual-based simulation of species range dynamics in a spatially varying environment. In brief, our model simulates a population invading a new environment and tracks population demography and the evolution of dispersal and local adaptation during range expansion.

#### The landscape

The simulation landscape consisted of a one-dimensional array containing 1000 patches. This setup is most akin to a natural landscape that is primarily one-dimensional, such as a river corridor, mountain ridge, or valley. The environment varies across space (*x*), producing a spatial gradient in phenotypic optima (θ) with slope *b*, which we vary across simulations (Figure 1a).

**Figure 1.**
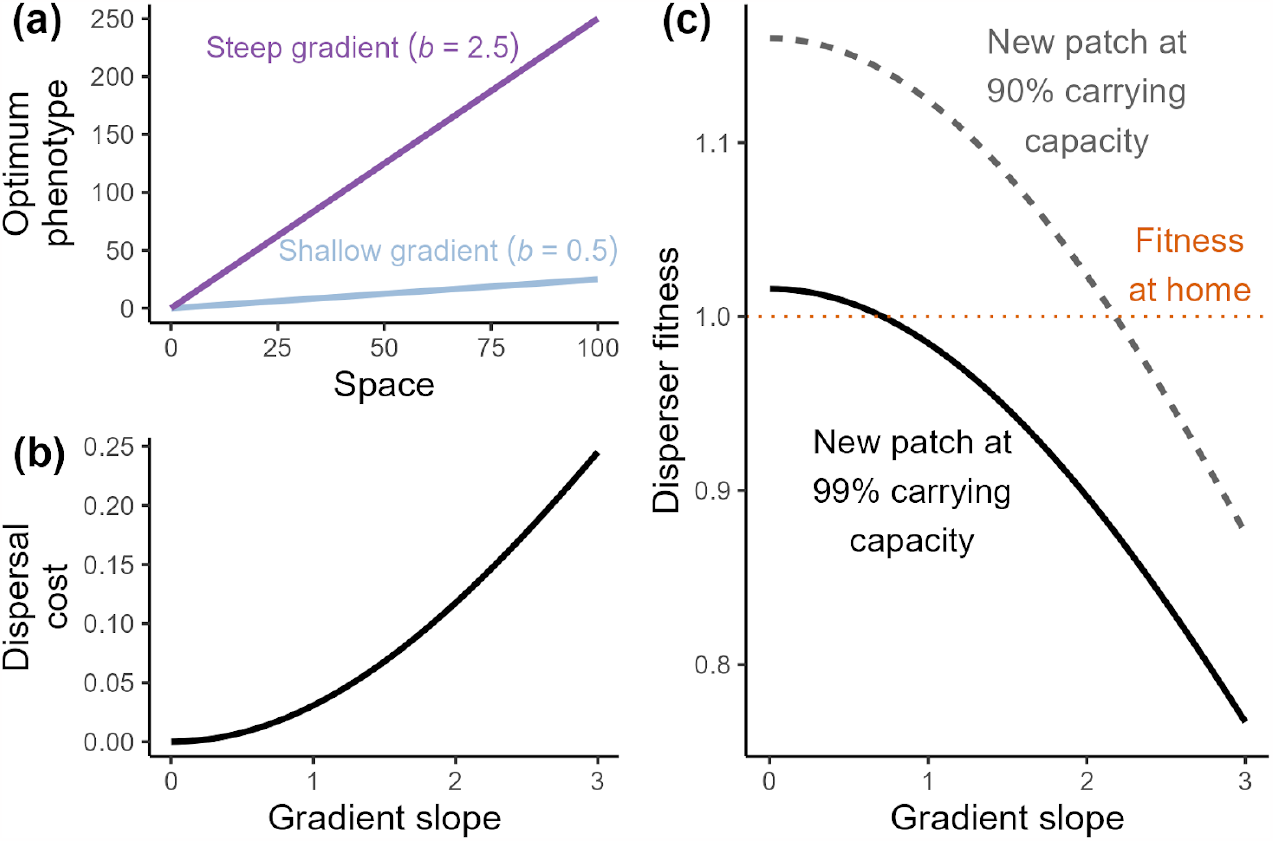
**(a)** Illustrates the simulation landscape (showing only 100 patches), where an underlying environmental gradient across space creates a gradient in optimal phenotype, whose slope can range from shallow (*b* = 0) to steep (*b* = 3). Examples of *b* = 0.5 and *b* = 2.5 are illustrated. All simulated invasions start at spatial position 0. **(b)** Shows the relationship between gradient slope (*b*) and dispersal cost, which is a function of gradient slope and the strength of selection. Dispersal cost is the decrease in P(survival) for an individual adapted to patch *x* dispersing to patch *x + 1*, calculated as 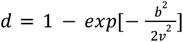, where *v* = 4. **(c)** Illustrates how the expected fitness of a dispersing individual (dispersing one patch away) will change based on the gradient slope (which influences viability fitness via maladaptation) and the population size in the new patch (which influences density dependent fecundity). Plot shows expected fitness at population densities of 99% (solid line) and 90% (dashed line) of carrying capacity. Fitness at home for a locally adapted individual will be 1, regardless of gradient slope (orange dotted line). Expected disperser fitness (y-axis) is the product of expected viability (*W*_*i*_) and expected fecundity (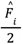 dividing by 2 as fecundity is calculated for females only; see Eqs. 1, 2 in Methods).

#### Genetics

In our simulations, individuals were diploid and either male or female, with obligate sexual reproduction, and ten chromosomes each 100,000 bp long. Generations were discrete. There were three mutation types: 1) neutral mutations, 2) mutations that contributed additively to a quantitative trait [i.e., biallelic quantitative trait loci (QTL) with no dominance] that determined fitness along the ecological gradient (niche trait), and 3) mutations that contributed additively to a quantitative trait determining dispersal propensity (dispersal trait). The overall mutation rate was set to 1 × 10^−7^ mutations per base position on a gamete per generation, and mutations were ten times more likely to be neutral than QTLs. Results were qualitatively similar with both lower and higher mutation rates (Fig. S1). The effect size for niche and dispersal QTLs were drawn from normal distributions (niche: N(0, 0.04), dispersal: N(0, 0.0004)). Mutation effect size distributions were different due to the difference in magnitude between these two traits (i.e., the niche trait might vary 0 - 100 across a landscape, but the dispersal trait was much smaller in magnitude to keep dispersal kernels realistic). Recombination rate was set to 1 × 10^−4^ (probability of a cross-over event between any two adjacent bases per genome per generation).

#### Mating, dispersal, and population dynamics

Individual fitness was a function of both survival and fecundity. Following (Bürger & Lynch 1995), viability fitness [P(*survival*)] for an individual (*W*_*i*_) was calculated as

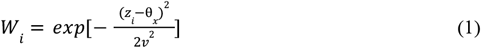

where θ_*x*_ is the phenotypic optimum in patch *x, v* is the strength of stabilizing selection (standard deviation of the fitness function; *v* = 4 throughout), and *z*_*i*_ is the individual’s niche trait phenotype. Thus the probability of survival is determined by the deviation of an individual’s phenotypic value from the local optimum in a Gaussian stabilizing selection model, with probability of survival decreasing with increasing maladaptation.

Following viability selection, individuals dispersed according to a Poisson dispersal kernel whose mean was defined by the individual’s dispersal trait phenotype, *m*_*i*_ (determined additively by the individual’s dispersal QTL). Dispersal direction (left or right along the gradient) was unbiased and random. Landscape boundaries were reflective such that individuals could not disperse outside of the first or last patch of the simulated landscapes.

Mating occurred after dispersal. Fecundity of female individuals was calculated as 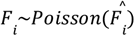, where

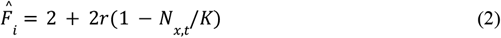

Thus, demographic stochasticity was included both through the survival process described above and the reproductive process described here. The maximum rate of increase (*r*) was set to 1.6. Fecundity was density dependent, based on the number of individuals in patch *x* at time t (*N*_*x,t*_) and the carrying capacity of each patch (here, *K* = 200 throughout). Male mates were drawn at random from the focal female’s patch for each offspring individually (i.e., multiple paternity was possible). Any 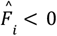 was set to 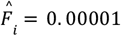, as the mean for a Poisson distribution in SLiM must exceed zero

For interpretability, in most of our results below we have transformed the spatial gradient slope, *b*, to represent the cost of dispersal due to maladaptation, which is the decrease in *P*(survival) for an individual adapted to patch *x* dispersing to patch *x + 1* (Fig. 1b). This dispersal cost is calculated as 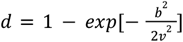. Representing the steepness of the environmental gradient as the expected decrease in survival increases its applicability and estimability in empirical work, as experiments could more easily measure survival in different locations along a gradient than estimate phenotypic optima along that gradient. The realized fitness of a dispersing individual will be a function of the dispersal cost in *P*(survival) due to maladaptation and density-dependent fecundity in the new patch (Fig. 1c).

### Simulation process

#### Burn-in

Each simulation started with a 20,000-generation burn-in phase. During this phase, 200 genetically uniform and perfectly adapted founding individuals were placed in the central patch of the landscape. Mating and dispersal occurred as described above, but the landscape was restricted to 11 patches (five on either side of the central patch). Every patch had a carrying capacity (*K*) of 200 individuals, resulting in a total landscape-wide carrying capacity of 2,200. A moderate spatial environmental gradient in optima spanned across the 11 patches (*b* = 0.5). In the presence of an environmental gradient, dispersal will be strongly selected against due to local adaptation (e.g., Holt 1985), unless there is some potential benefit of dispersal (e.g., colonization of a patch with low competition due to recent patch extinction). Thus, to maintain genetic variation for dispersal during the burnin, one patch was randomly selected to go extinct every other generation. At the 20,000th generation, all individuals in the central patch migrated to the first patch of the main landscape (i.e., the “founding event”); these two patches had the same phenotypic optimum. All other individuals were removed from the simulation.

#### Main simulation

Following the burn-in phase, the main simulation commenced using the specified parameters (Table S1). The landscape was expanded to 1000 patches, with the first (left-most) patch containing the founding individuals described above. The slope in phenotypic optima across the landscape was determined by *b*. Similar to extinction during the burnin, to prevent dispersal from evolving to zero in occupied portions of the range due to persistent selection against dispersal, ten patches (one percent of the landscape) were randomly selected for extinction each generation (regardless of whether the patch was colonized). Results were largely insensitive to differing extinction rates (Fig. S2). The simulation ended after 200 generations, or if all populations became extinct, or if the furthest landscape patch achieved a population size of at least half the carrying capacity (i.e., the species occupied the entire landscape). We conducted 1000 simulations with values of *b* randomly drawn from a uniform distribution: *b∼U*(0, 3). To compare our results to simulations with no dispersal evolution, we ran another 1000 simulations where dispersal was static and did not evolve [*m* = 0.55, which was the mean dispersal of the founding populations (after burnin) in the simulations with dispersal evolution]. We also ran sensitivity analyses to test the effects of 1) carrying capacity, selection strength, and intrinsic growth rate, 2) background extinction rate, and 3) mutation rate at niche and dispersal QTL, on model outcomes (SI *Sensitivity analyses*).

## Results

### Environmental gradients oppose the evolution of increased dispersal

As environmental gradients steepen, the change in phenotypic optimum between patch *x* and patch *x + 1* increases, increasing the fitness cost of dispersal (Fig. 1). Thus, environmental gradients oppose the evolution of higher dispersal at the invasion front as this dispersal cost opposes the evolution of increased dispersal (Fig. 2). For interpretability, we transform the environmental gradient slope (x-axis in Fig. 2 a,b) into the dispersal cost of maladaptation (x-axis in Fig. 2c), which is the decrease in P(survival) for an individual adapted to patch *x* dispersing to patch *x + 1* (see Methods; Fig. 1b). Dispersal evolution at the invasion front decreases sharply as dispersal cost increases from 0 (homogenous environment) to ca. 0.01, then more gradually until a dispersal cost of 0.1, after which there is no signal of dispersal differentiation between the range edge and core (Fig. 2c). These results are quantitatively similar across a wide range of carrying capacities, selection strengths, and intrinsic growth rates (Fig. S3).

**Figure 2.**
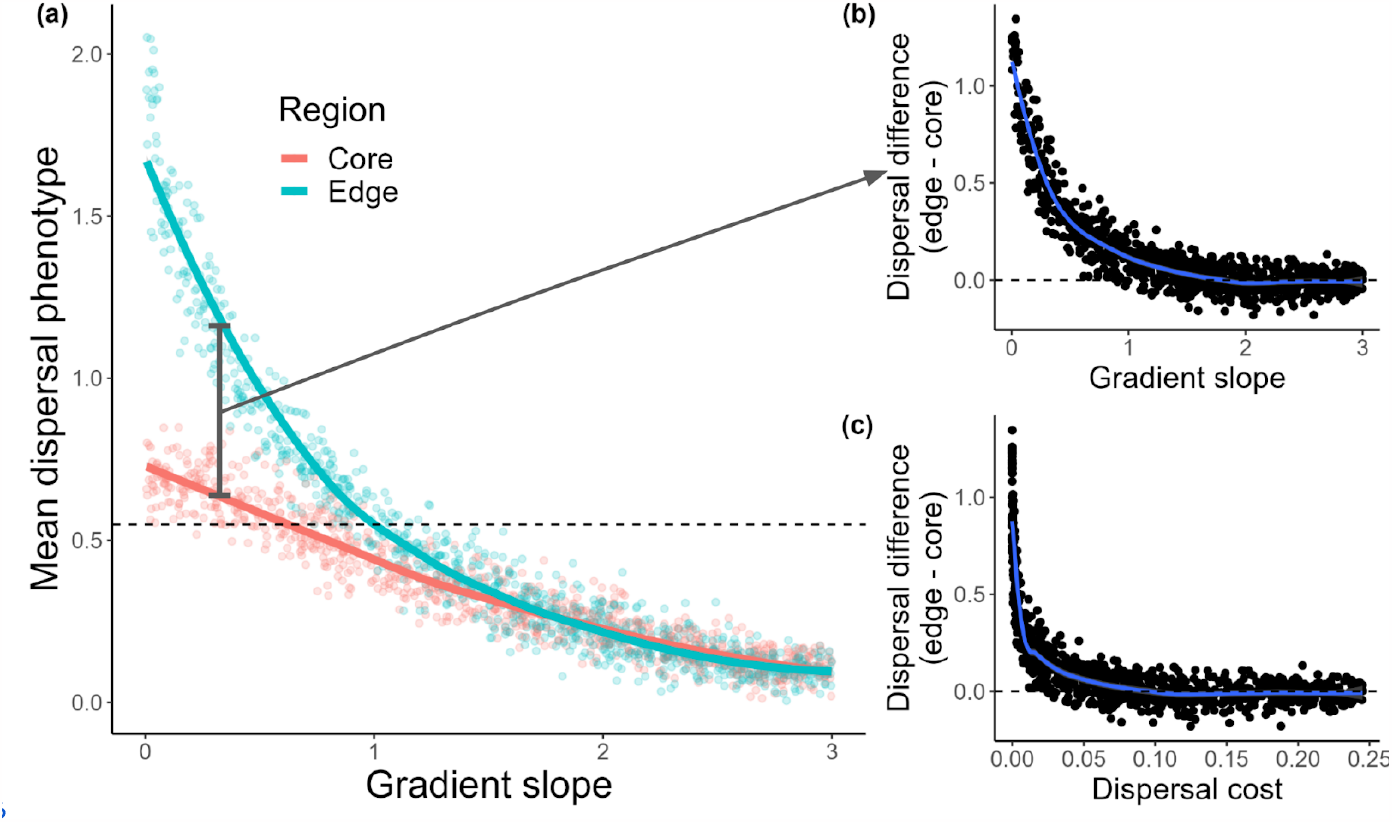
Environmental gradients oppose the evolution of increased dispersal during range expansion. Results of 1000 simulations after 200 generations of expansion. **(a)** Illustrates the mean dispersal phenotype within core and edge regions plotted against spatial gradient slope. Core and edge regions comprise the five most proximate and distal patches to the founding patch, respectively. The dashed line signifies the mean dispersal of the founding populations at the start of expansion. **(b)** Depicts the difference in dispersal between the core and edge across gradient slopes, expressed as the absolute increase in mean dispersal ability of the edge relative to the core. To aid in interpretability, **(c)** shows the magnitude of dispersal evolution plotted against dispersal cost instead of gradient slope. Dispersal cost is a function of gradient slope (*b*) and the strength of selection (*v*), and is the expected decrease in *P*(survival) for an individual perfectly adapted to patch *x* dispersing to patch *x+1* (Methods).

### Environmental gradients make invasion slower and more predictable, especially when dispersal can evolve

Dispersal evolution greatly increased invasion distance relative to models with static dispersal, but this phenomenon was most apparent along shallow environmental gradients where dispersal cost is low (Fig. 3a). As dispersal cost increased beyond ∼ 0.01 (i.e., a 0.01 decrease in P(survival) for individuals dispersing from patch *x* to patch *x+1*), the difference between scenarios with and without dispersal evolution was comparatively trivial in magnitude. Surprisingly, evolution of decreased dispersal along steep gradients (dispersal cost ≳ 0.1) led to greater rates of expansion compared to scenarios with no dispersal evolution, as lower dispersal allowed better adaptation, and thus higher intrinsic growth rate. The increased unpredictability of invasion due to dispersal evolution found in earlier work (e.g., Ochocki & Miller 2017) was largely restricted to very shallow environmental gradients (Fig. 3b). The evolution of lower dispersal across steep gradients tended to increase predictability (decreased CV) compared to scenarios without dispersal evolution, though across very steep gradients CV’s for the two scenarios were similar (Fig. 3b) .

**Figure 3.**
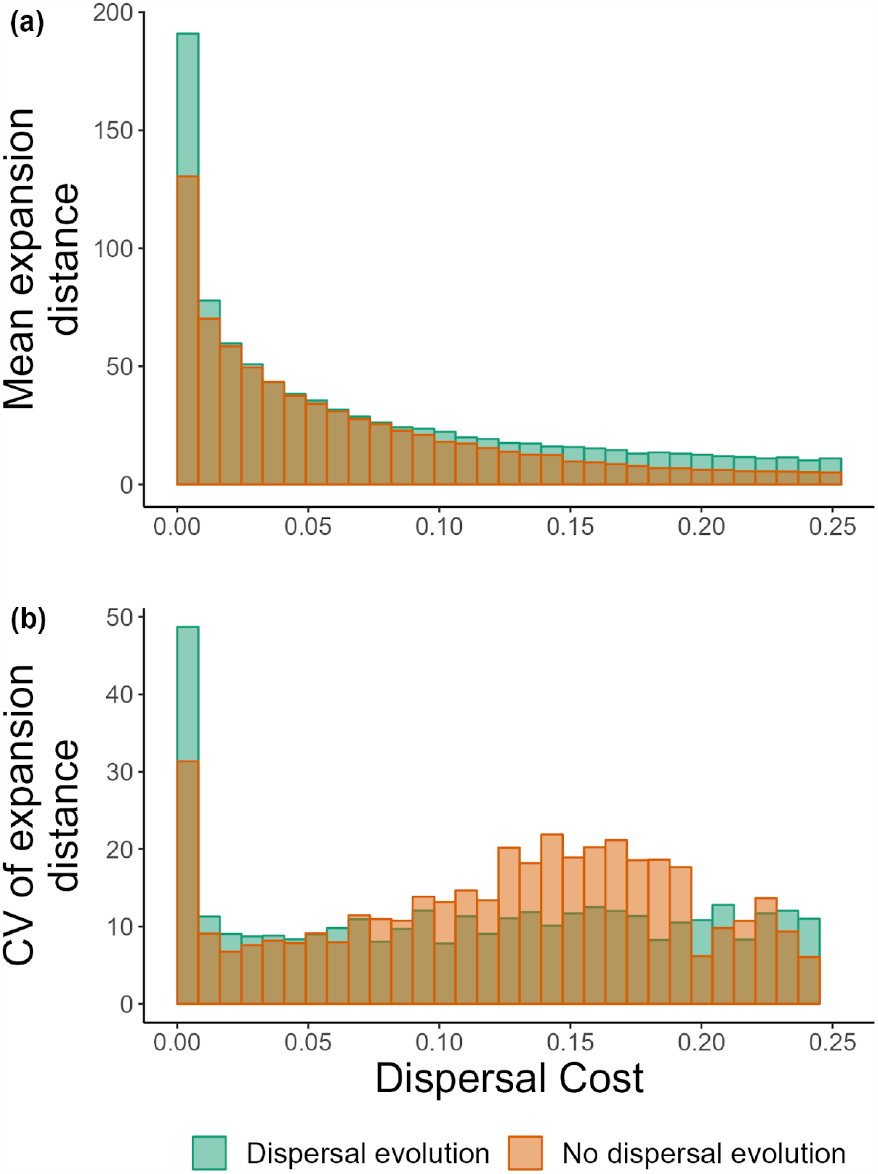
Environmental gradients make invasion slower and more predictable, especially when dispersal can evolve. **(a)** A binned summary plot showing the mean expansion distance within bins of dispersal cost for scenarios with (green) and without (orange) dispersal evolution. **(b)** A binned summary plot showing the coefficient of variation (CV) of expansion distance within bins of dispersal cost, again for scenarios with (green) and without (orange) dispersal evolution.

### Evolution of decreased dispersal along steep gradients saves populations from maladaptive gene flow

Increased dispersal leads to higher gene flow, which generally increases genetic variance within populations. Thus, along shallow environmental gradients in our simulations, the evolution of increased dispersal tended to increase the amount of genetic variance in the niche trait, relative to scenarios without dispersal evolution (Fig. 4a). However, as environmental gradients steepened and dispersal became more costly, dispersal evolved downward, as selection against dispersal became stronger (Fig. 2a). Thus, after dispersal evolution switched from increased to decreased dispersal (roughly at the dispersal cost marked with a vertical dashed line in Fig. 4), the evolution of decreased dispersal tended to lower the amount of genetic variance in edge populations relative to models with static dispersal. Lower genetic variance in the niche trait across steep gradients tended to increase fitness in edge populations (Fig. 4b,c) because populations experienced lower genetic load (i.e., fewer individuals are far from the patch phenotypic optimum). This decreased load is why dispersal evolution increased expansion distance relative to scenarios without dispersal evolution along steep gradients (Fig. 3a).

**Figure 4.**
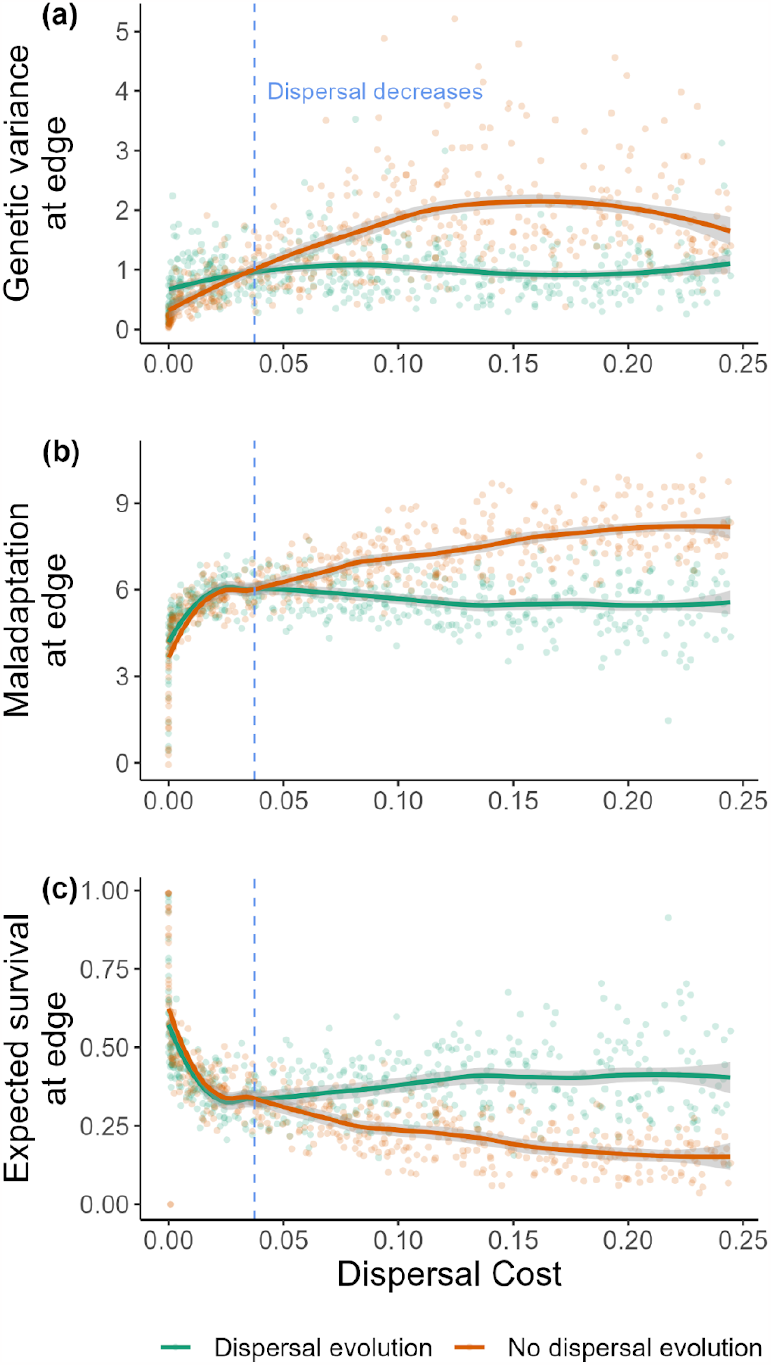
Evolution of decreased dispersal along steep gradients saves populations from maladaptive gene flow and increases expansion speed. **(a)** Genetic variance for the niche trait at the invasion front for simulations varying in dispersal cost (x-axis), after 200 generations of invasion, for scenarios with (green) and without (orange) dispersal evolution. **(b)** Maladaptation (deviation of mean niche trait from optimum) of edge population. **(c)** Represents the expected survival at the invasion front for the same scenarios and conditions. Expected probability of survival is calculated using the mean and variance of the population’s niche trait, the optimal niche trait value, and the selection strength (SI “Expected survival”). In all panels, the edge population in each simulation is the most distal occupied patch with at least ten individuals. Colored lines in each panel are loess regressions with 95% confidence bands. The blue vertical dashed line in all panels marks the approximate dispersal cost at which dispersal in the edge population begins to evolve downward (i.e., is lower than the dispersal of the initial founding population), estimated from the data shown in Fig. 2a.

### Spatial patterns of dispersal are unstable through time

When the landscape is homogenous (i.e., no fitness cost of dispersal), the evolution of increased dispersal at the invasion front was permanent and the spatial pattern of dispersal phenotypes was constant through time (Fig. 5a). However, as dispersal cost increased, the spatial pattern of dispersal phenotypes across the landscape was increasingly unstable through time (Fig. 5b-d). Though spatial sorting and density dependence served to increase dispersal at the invasion front, over time, populations at the invasion front tended to evolve lower dispersal as selection acted against dispersal. Importantly, spatial sorting and density dependence acted in a spatially explicit manner to create clines in dispersal phenotypes through space while selection against dispersal due to steep gradients affected the entire landscape equally. This means that while landscape-wide selection against dispersal lowered average dispersal phenotypes through time, the spatial cline in relative phenotypes from core to edge was still observed even after many generations.

**Figure 5.**
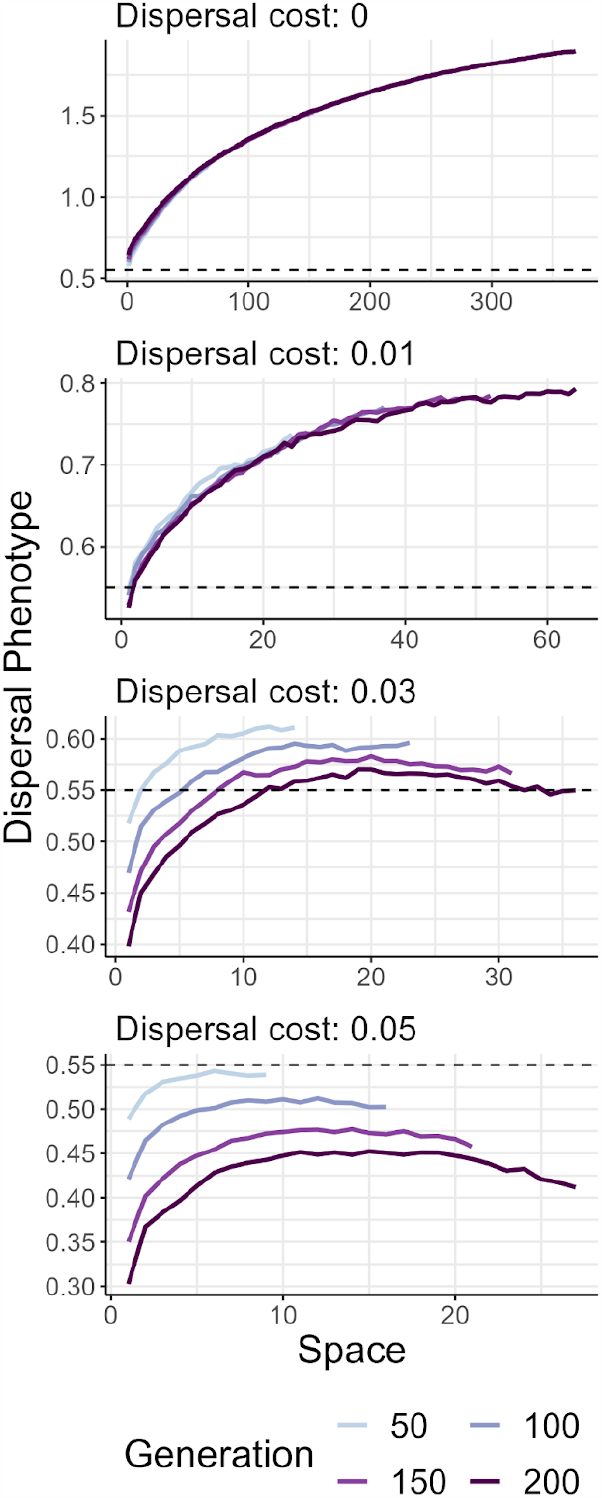
Spatial patterns of dispersal are increasingly unstable through time as environmental gradients steepen. Mean dispersal phenotypes are plotted for each occupied patch along the landscape, at different timepoints of invasion (line color). Lines show means across 40 simulations for each dispersal cost. Panels show increasingly steep environmental gradients, from a homogenous landscape (dispersal cost: 0) to a moderately steep landscape (dispersal cost: 0.05). A dashed horizontal line indicates the mean dispersal phenotype of the founding populations, averaged across simulations.

Local adaptation and dispersal evolution during invasion — the *Diorhabda* range expansion

To illustrate how our model could be paired with empirical data to predict invasions and inform management decisions, we parameterized the model with data from a recent range expansion of a biocontrol agent. *Diorhabda carinulata*, the northern tamarisk beetle, was released in 2001 into the western United States for the biological control of invasive riparian shrubs in the genus Tamarix, or saltcedar or tamarisk (DeLoach et al., 2003). The range of the beetle was initially limited to areas north of 38°N, because it was maladapted to the photoperiods in the south, which are shorter midsummer than northern daylengths, and initiated diapause (seasonal dormancy in insects, similar to hibernation in mammals) too early in the season (Bean *et al*. 2007). Because both photoperiod and the timing of winter onset change with latitude, there is a gradient in the optimal photoperiod cue to initiate diapause from north to south across the range of the beetle. Nearly a decade after the first releases, the beetles started to disperse southward following remote riparian corridors, enabled by evolution of the photoperiod cue that initiates diapause (Bean et al., 2012). In 2018, Clark et al. (2022) measured dispersal traits using tethered flight mills for beetles from the core of the range and the leading edge of the expansion, reared in a common lab environment, and found that dispersal propensity and ability had increased in beetles from the leading edge. Also in 2018, Clark et al. (2023) determined that the photoperiod cue for diapause had shifted in edge beetles compared to core beetles, such that some populations from the core and edge had become locally adapted to photoperiod. The tamarisk beetle system, showing both evolution of dispersal and local adaptation to an environmental gradient during range expansion, provides a useful system with which to parameterize our model.

We used Approximate Bayesian Computation (ABC) to estimate the gradient slope and strength of selection (which together describe the dispersal cost) during the tamarisk beetle range expansion. ABC is a statistical method used to estimate model parameters when likelihood calculations are intractable, by comparing simulated to observed results via a set of summary statistics (reviewed in Beaumont 2010). We calculated three summary statistics: the difference in dispersal phenotype between edge and core populations, the difference in niche phenotype between edge and core populations, and the ratio in survival between edge and core populations in the core habitat (a measure reflecting local adaptation and the strength of selection). The observed summary statistics were calculated using data from experiments by Clark et al. (2022, 2023), where the dispersal phenotype was the mean distance flown in individual dispersal trials, the niche phenotype was the days until diapause, and survival was the proportion of individuals in diapause after the diapause cue was initiated. We used data from two tamarisk beetle populations that represented core and edge populations after range expansion ∼500 km southward across ∼34 generations. To estimate posterior distributions for gradient slope (*b*) and the strength of selection (*v*), we ran 60,000 simulations, sampling each parameter from a uniform prior [*b∼U*(0, 2); *v∼U*(0, 5)]. We estimated a joint posterior of dispersal cost, *d*, based on 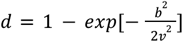. Full details of analyses are in **SI: ABC analysis**.

ABC estimated dispersal cost during the southward tamarisk beetle expansion as a decrease in *P*(survival) of ∼ 0.025 per 36 km (Fig. B1a). If there were *not* a gradient in optimal phenotype that introduced a dispersal cost (i.e., if the tamarisk beetle was expanding across a homogenous landscape), simulations suggested that expansion could have reached ∼1700 km, as opposed to the observed ∼500 km. Thus, local selection along a latitudinal gradient has likely slowed the tamarisk beetle expansion considerably. Simulations suggested this slowdown was attributable to both the direct negative effect of maladaptation on colonization of new habitat, and the constraint that dispersal cost placed on dispersal evolution at the invasion front (Fig. B1b). These analyses demonstrate how models such as ours can be paired with empirical data to better understand and predict the course of invasion, while extensions of our model could be used to model effectiveness of different management interventions for slowing invasions.

**Figure B1.**
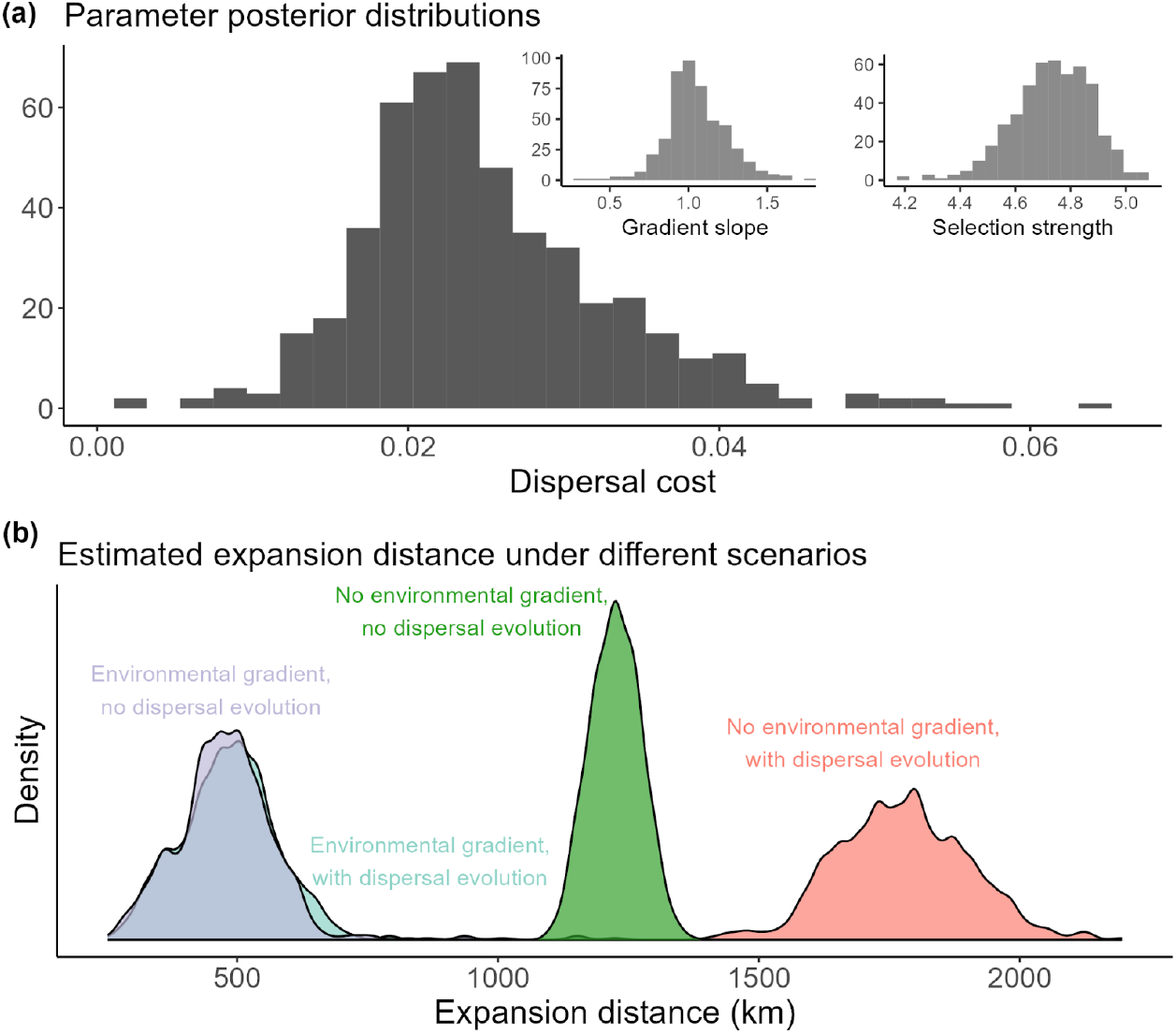
**(a)** Using ABC to integrate our model with empirical data on *Diorhabda* expansion, we estimated a posterior distribution for the joint parameter, dispersal cost (*d*), which is calculated based on gradient slope (*b*) and selection strength (*v*) (insets). **(b)** Comparing expansion distances for simulations using the estimated posteriors for *b* and *v* (“Environmental gradient, with dispersal evolution); assuming the estimated gradient but with no dispersal evolution (“Environmental gradient, no dispersal evolution”); assuming a homogenous gradient where *b* = 0 (“No environmental gradient, with dispersal evolution”); and assuming a homogenous gradient with no dispersal evolution (“No environmental gradient, no dispersal evolution”); see **SI: ABC analysis** for full details.

## Discussion

Populations invading across a landscape may be subject to evolutionary forces that manifest on ecological timescales to alter the trajectory of invasion. Much work has focused on how environmental heterogeneity can result in local selection influencing invasion (e.g., Kirkpatrick & Barton 1997; Polechová 2018), and how the evolution of increased dispersal at the expansion front can accelerate invasions (e.g., Travis & Dytham 2002; Phillips *et al*. 2006; Shine *et al*. 2011). However, we have a limited understanding of how these two forces interact during invasion. Given that expansion speed is a function of intrinsic growth rate and dispersal ability (Hastings *et al*. 2004), exploring how environmental heterogeneity influences fitness and dispersal evolution is crucial to understanding species’ range expansions. Here, we showed how environmental gradients oppose the evolution of increased dispersal during invasion, leading to a strong signal of dispersal evolution only along shallow environmental gradients. We found that while evolution across homogenous landscapes can cause invasions to be highly unpredictable, spatially varying selection across even shallow environmental gradients greatly increased the predictability of invasion in our simulations. Along steep gradients, the evolution of lower dispersal allowed increased adaptation and thus increased population growth rates, which enhanced invasion speed relative to scenarios without dispersal evolution. Though increased dispersal at the invasion front could persist for hundreds of generations, temporal shifts in the relative strength of dispersal evolution and local selection led to population mean dispersal being unstable through time. Overall, we find that local adaptation and dispersal evolution are deeply entwined, with potentially large influences on our understanding of invasion.

### The interplay between local adaptation, dispersal cost, and dispersal evolution

Local adaptation within spatially heterogeneous environments is common in both native and invasive species (Hereford 2009; Oduor *et al*. 2016; Briscoe Runquist *et al*. 2020). We can also expect many populations to harbor genetic variation in dispersal traits (reviewed in Ronce 2007). Given these two conditions — spatially varying selection and heritable dispersal ability — the processes of local adaptation and dispersal evolution will interact. As environmental gradients steepen and phenotypic optima become more disparate across space, the fitness cost of dispersal increases, and thus maladaptation will oppose the evolution of increased dispersal. Consequently, environmental gradients reduce invasion speed and increase the predictability of invasion.

There are now multiple observations that support theoretical predictions of increased dispersal ability at the edge of an invading species range (Simmons & Thomas 2004; Phillips *et al*. 2006; Weiss-Lehman *et al*. 2017). However, the phenomenon is not ubiquitous (Miller *et al*. 2020; Oldfather *et al*. 2021), and predictions for the trajectory of dispersal evolution during invasion remain elusive. Prior theory has indicated that the evolution of higher dispersal at the invasion front can be constrained by tradeoffs between dispersal and other traits (Ochocki *et al*. 2020), interspecific competition (Burton *et al*. 2010), strong Allee effects (Travis & Dytham 2002), and availability of suitable habitat (Travis & Dytham 1999). Our model suggests that dispersal evolution will be strongly mediated by the steepness of the environmental gradient a species is invading across. In general, large increases in the rate and unpredictability of invasions due to dispersal evolution should be limited to cases where environmental gradients are relatively shallow and the cost of dispersal is low.

In accord with recent empirical work (Ochocki & Miller 2017; Weiss-Lehman *et al*. 2017), our model showed that when landscapes are homogenous, evolutionary processes acting on dispersal traits will generally decrease the predictability of invasion across instances (Fig. 3b). However, environmental gradients greatly increased the predictability of invasion in our model, regardless of whether dispersal could evolve or not. In the absence of a gradient, individuals in our simulation could conceivably survive and reproduce in any location throughout the landscape. As dispersal in our model was stochastic, even low dispersal trait values could result in low-probability long-distance dispersal events and, without a gradient, these long-distance events often led to successful establishment. Thus, variation in expansion distances was largely driven by these stochastic long-distance dispersal events in homogeneous landscapes. However, spatially varying selection along even shallow gradients changed this dynamic by imposing a fitness cost to dispersal (i.e., maladaptation). Unlike other fitness costs sometimes modeled with dispersal (e.g., a constant risk of mortality), the cost imposed by a gradient will increase with distance such that it is greatest for long-distance dispersal events. Thus, even shallow gradients restricted the successful establishment of rare, long-distance dispersal events, reducing stochasticity in invasion speed across instances.

### Evolution of *decreased* dispersal may be important for structuring species distributions

While most work addressing dispersal evolution during range expansion has focused on increases in dispersal ability, our results suggest intriguing possibilities for the evolution of decreased dispersal. When the fitness cost of dispersal is greater than the benefit of escaping competition, there will be selection against dispersal. This will be most likely in scenarios with some combination of steep gradients, high initial dispersal, and/or strong local selection. During range expansion, somewhat counterintuitively, the evolution of lower dispersal along steep gradients allows for increased expansion rates (Fig. 3a). This is because lower dispersal reduces maladaptive gene flow, which allows local adaptation to increase the population growth rate (Fig. 4). This result emphasizes the fact that expansion speed is determined by the combination of dispersal and population growth rate — when reduced dispersal increases the growth rate via local adaptation, this can positively impact expected speed. Furthermore, evolution of lower dispersal during expansion across steep gradients increased the predictability of invasion compared to scenarios with static dispersal (Fig. 3b), as decreased gene flow and higher local adaptation resulted in less stochastic population dynamics. These insights have potentially important consequences for our understanding of the “critical environmental gradient” (a gradient so steep that range expansion is wholly prevented) in earlier range limit models with static dispersal (e.g., Polechová & Barton 2015; Polechová 2018). Populations may be able to spread across steeper gradients than expected if the evolution of lower dispersal rates facilitates local adaptation.

### Spatial patterns of dispersal will often change through time

In our model, the spatial pattern of dispersal was unstable as the net effects of dispersal, density dependence, and local adaptation shifted through time. At the invasion front, spatial sorting and selection will often increase dispersal, but if there is a fitness cost to dispersal, this will be opposed by maladaptation. We will likely often see an initial transitory phase of invasion at the expansion front, where dispersal increases rapidly due to spatial sorting and low population size. After the invasion wave passes, these populations will exhibit more stable dynamics dominated by selection, intraspecific competition, and the evolution of decreased dispersal (e.g., Travis & Dytham 2002; Simmons & Thomas 2004; Clarke *et al*. 2019). While dispersal did decline through time in the presence of environmental gradients, heightened dispersal at the edge relative to the core persisted for hundreds of generations in our simulations, meaning that when it occurs, dispersal evolution will likely be important on the timescales relevant to management of native and invasive populations.

### Predicting and managing range expansions

What do these results mean for range expansion of natural populations? First, we can expect that elevational gradients will be steeper than latitudinal gradients (e.g., Bachmann *et al*. 2020). Thus we may be less likely to see evolution of increased dispersal during invasion along elevational gradients. Of course, for native species where persistence will require upslope range expansion (Geppert *et al*. 2023), such local selection pressure could hinder the potentially positive effects of dispersal evolution in accelerating tracking of suitable habitat (Weiss-Lehman & Shaw 2020). Second, our work suggests that while invasions are surely hard to predict, the most pessimistic hurdles of unpredictability should only manifest across near-homogenous gradients. This means that, given the right data, predicting invasion dynamics may be more feasible than it would otherwise seem.

Of course, in practice, we only rarely have the data needed to parameterize such predictive models — namely, dispersal ability, growth rate, and gradient steepness. But these data are straightforward, if not easy, to collect. A long history of common garden and transplant studies have assessed fitness differences among populations sourced at varying distances from the planting site (Antonovics & Bradshaw 1970; Lee *et al*. 2003; Maron *et al*. 2004; Colautti & Barrett 2013; Bachmann & Van Buskirk 2021; Gorton *et al*. 2022; Peschel & Shaw 2023), which can be used to calculate fitness cost per unit distance (i.e., how quickly is fitness expected to decline for an individual dispersing from its natal patch). Dispersal rates can be estimated based on spread rates or dispersal trials (e.g., Clark *et al*. 2022). Parameterizing models like ours with such data could be a fruitful tool for the management of invasions. In general, recognition of the interplay between dispersal and local adaptation, as well as the temporal instability of dispersal evolution, will lead to a greater understanding of species range expansions and more predictive models of spread dynamics.

## Acknowledgements

We thank the Weiss-Lehman lab for comments on an earlier draft of this manuscript. This work was supported by grants from the National Science Foundation (DBI-2010892 to JWB, EPS-2019528 to CW, DEB-2230806 to CW and JWB, and DEB-2121980 to RAH.) and the United States Department of Agriculture (NIFA AFRI Predoctoral Fellowship 2021-09368 to EIC). Any opinions, findings and conclusions expressed in this material are those of the authors and do not necessarily reflect the views of NSF or USDA.

