## Supplementary material for "Environmental gradients mediate dispersal evolution during biological invasions": SI

### Sensitivity analyses

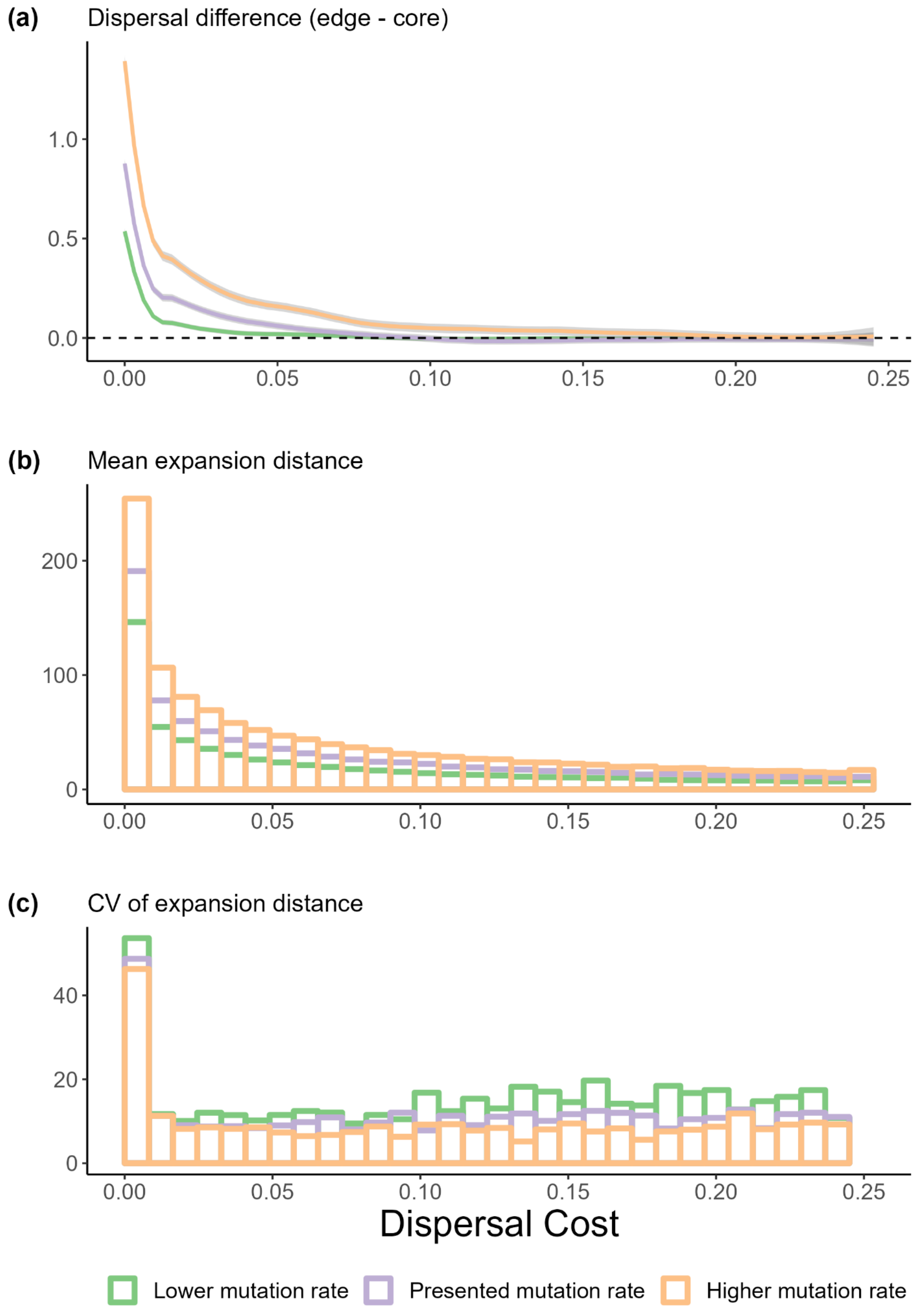


**Figure S1.** Comparing results for scenarios with presented ($1 \times 10^{-8}$, used in the main simulations; purple), lower ($5 \times 10^{-9}$; green), and higher ($2 \times 10^{-8}$; orange) mutation rates for the niche and dispersal QTLs. **(a)** depicts the difference in dispersal between the core and edge across gradient slopes (expressed as the absolute increase in mean dispersal ability of the edge relative to the core) **(b)** A binned summary plot summarizing the mean expansion distance within bins of dispersal cost **(c)** A binned summary plot summarizing the coefficient of variation (CV) of expansion distance within bins of dispersal cost. Compare to Figures. 2, 3 in the main text.

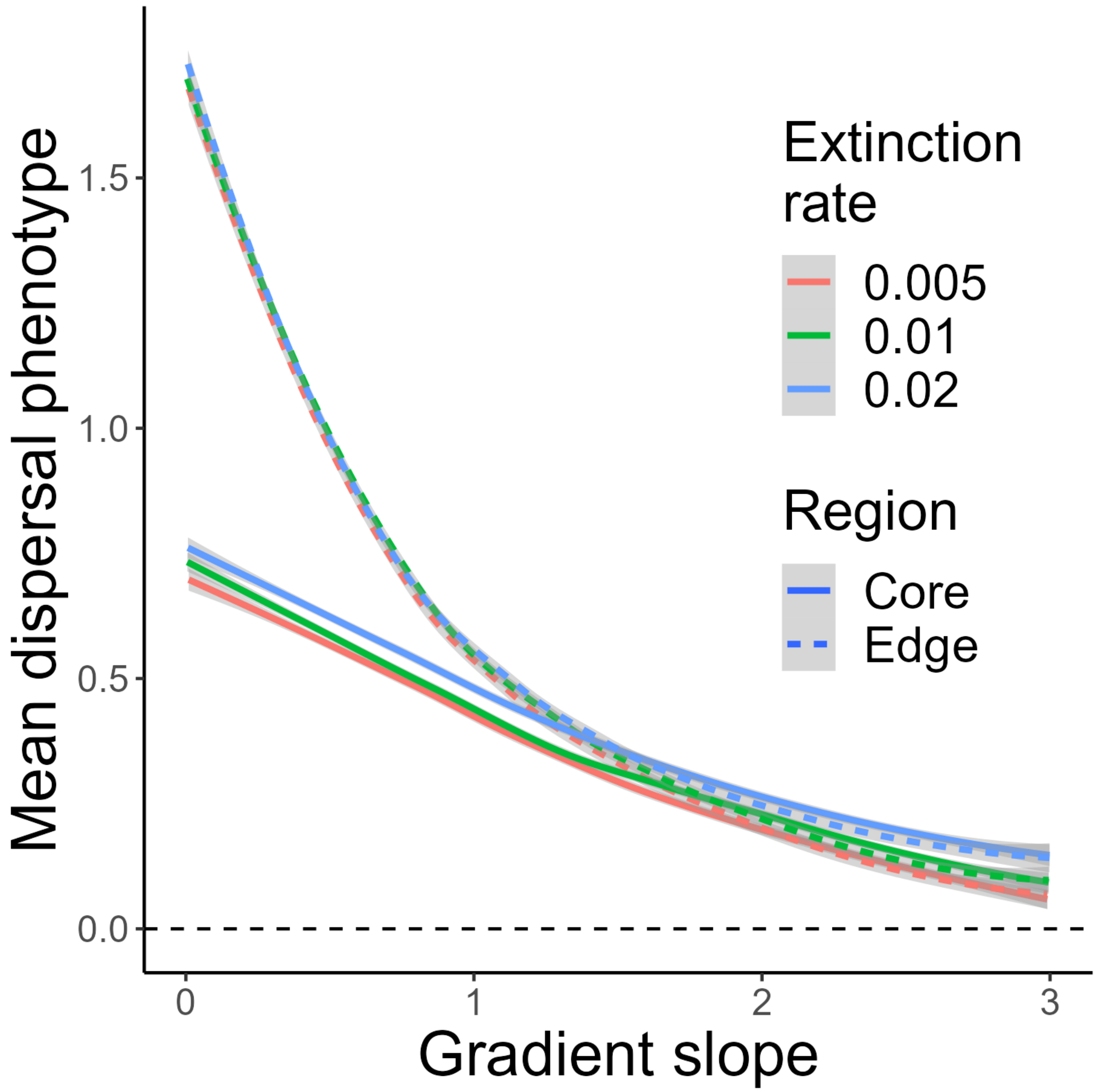


**Figure S2.** Patterns of dispersal evolution in core vs. edge regions with varying patch extinction rates. Compare to Fig. 2a in the main text. An extinction rate of 0.01 was used in the main simulations.

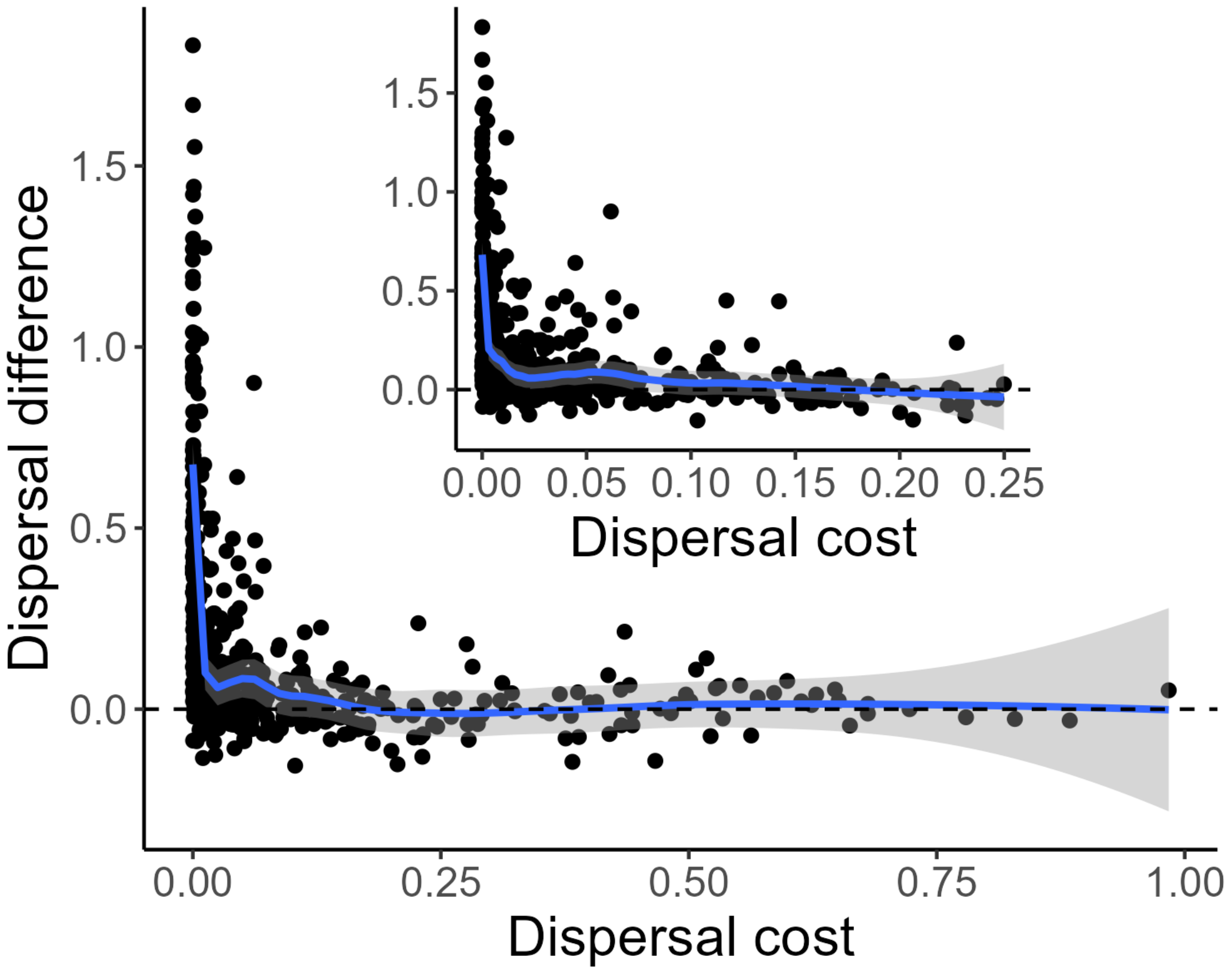


**Figure S3.** Patterns of dispersal evolution with changing dispersal cost, across varying patch carrying capacities (K), selection strengths (v), and intrinsic growth rates (r). Compare to Fig. 2c in main text. *b* [U(0,3)]; *v* [U(0, 10)]; *K* / *K_b* [U(50, 300)]; *r* [U(0.5, 2.5)]. Inset restricts x-axis range to be comparable to Fig. 2c to aid in comparison.

Supplementary Information for: *Environmental gradients mediate dispersal evolution during biological invasions*

### Tables

Table S1. Parameter values for the SLiM simulation model.

| Parameter | Definition | Value |
|---|---|---|
| *Varying parameters* | | |
| $b$ | Slope of spatial gradient in phenotypic optima | U(0-3) |
| *Static parameters* | | |
| $K$ | Patch carrying capacity (individuals) | 200 |
| $r$ | Maximum rate of increase | 1.6 |
| $v$ | Strength of stabilizing selection (SD of fitness function) | 4 |
| $m$ | Initial expected dispersal per generation (mean of Poisson dispersal kernel) at beginning of burn-in | 0.5 |
| $e$ | Patch extinction rate | 0.01 |
| $\mu$ | Mutation rate per base position per generation | $10^{-7}$ |
| ρ | Recombination rate (crossover events per base position per generation) | $10^{-4}$ |

### ABC analysis

We used Approximate Bayesian Computation (ABC) via the abc R package (Csilléry *et al.* 2012) to parameterize our model using empirical data from the range expansion of the tamarisk beetle (*Diorhabda carinulata*) in the Western United States (see **Box 1** in main text). ABC is a statistical method used to estimate model parameters when likelihood calculations are intractable (reviewed in Beaumont 2010). In brief, ABC operates in the following manner: initially, numerous simulations are executed with parameter values drawn from defined prior distributions. Next, for each simulation, summary statistics (measures that describe relevant features of the data) are computed and compared with those from observed data. This comparison yields a distance metric for each simulation, indicating how much the simulated and actual summary statistics differ. Users set a tolerance level to determine which simulations to keep (for example, a 0.01 tolerance level means keeping the top 1% of simulations with the smallest distance metrics). The parameter values from these selected simulations are then considered samples from the posterior distribution of the parameter, with various methods available to construct this distribution. To assess the model's adequacy, graphical posterior predictive checks are commonly used. This involves generating another set of simulations using parameters from the estimated posterior distributions, and then comparing the distribution of their summary statistics against the observed summary statistics. If the observed statistics are significantly different or at the extreme ends of the simulation results, it suggests that the model may be inadequate.

For our ABC analysis, we calculated three summary statistics: the difference in dispersal phenotype between edge and core populations, the difference in niche phenotype between edge and core populations, and the ratio in survival between edge and core populations in the core habitat (a measure reflecting local adaptation and the strength of selection). For each simulation, these statistics were calculated for core and edge using dispersal and niche phenotype values averaged over the three patches at each end of the distribution after 34 generations of spread. The observed summary statistics were calculated using data from lab dispersal and common garden experiments by Clark et al. (2022, 2023), using two tamarisk beetle populations that represented core and edge populations after range expansion ~500 km southward across ~34 generations. See Clark et al. (2022, 2023) for full experimental details; in brief,

- dispersal phenotype was the population average for distance in meters a male beetle flew during a 1 hour dispersal trial
- the niche phenotype was the population average for number of days a female took to enter diapause at a constant photoperiod that would typically cause 50% of a northern (core) population to diapause
- survival was the population average for proportion of females in diapause after 42 days at a constant photoperiod that would typically cause 50% of a northern (core) population to diapause (with the assumption that those individuals not entering diapause would not survive)

So that summary statistics could be readily compared between simulated and observed data, dispersal and niche phenotype measures were scaled (centered at zero with $\sigma = 1$) across simulation results, and separately across the eight tamarisk beetle populations measured in Clark et al. (2022, 2023), before calculation of summary statistics.

To estimate posterior distributions for gradient slope (*b*) and the strength of selection (*v*), we ran 60,000 simulations for 34 generations of expansion, sampling each parameter from a uniform prior $[b \sim U(0, 2);\ v \sim U(0, 5)]$. Other model parameters were fixed (see values in Table 1). Parameter posterior distributions for *b* and *v* were estimated using localized linear regression and a tolerance rate of 0.01. We estimated a joint posterior of dispersal cost, *d*, based on $d = 1 - exp[-\frac{b^2}{2v^2}]$. Graphical posterior predictive checks were performed using 500 simulations run with values of *b* and *v* drawn from their estimated parameter posterior distributions. Observed summary statistics fell well within the distributions of these posterior predictive checks (**Fig. S4**).

To express the estimated dispersal cost in terms of the decrease in $P$(survival) across geographic distance, as opposed to "per patch", we assumed that the mean expansion distance in our posterior predictive check simulations (14 patches) represented the expansion distance of the actual tamarisk beetle expansion (~500 km), such that distance between patches corresponded to 500 / 14 = 36 km. To estimate how much the presence of an environmental gradient constrained range expansion, we ran another 500 simulations, using the same $v$ values as the posterior predictive checks, but with a homogenous gradient ($b = 0$). To estimate how much dispersal evolution during expansion contributed to increased expansion, we compared those simulation results to two other batches of simulations (across the estimated gradient, and across a homogenous gradient), but where dispersal was static (i.e., not evolving).

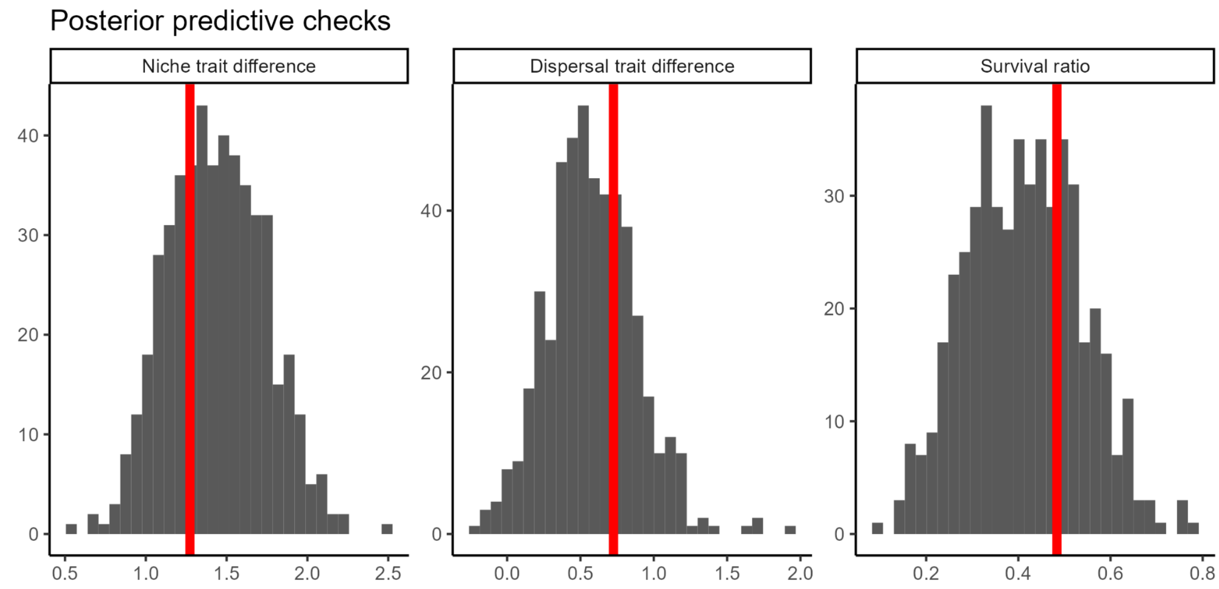


**Figure S4.** Posterior predictive checks of the three summary statistics. Vertical red lines mark the observed summary statistics. Trait differences are the differences between the (scaled) trait values of the edge and core populations; survival ratio is the ratio in survival between edge and core populations in the core habitat.

### Expected survival

The expected mean *P*(survival) of a population shown in Fig. 4c was calculated based on the optimal trait value in that population's patch (θ), the population mean trait value (μ), the standard deviation of the trait distribution (σ), and the strength of stabilizing selection ($v$). Numerical integration was used to find the integral,

$$E[survival] = \int_{\theta-100}^{\theta+100} w(z) \bullet f(z)\, dz,$$

where $w(z) = exp(-\frac{(z-\theta)^2}{2v^2})$ is the fitness function (Equation 1 in the main text) and

$f(z) = \frac{1}{\sigma\sqrt{2\pi}} exp(-\frac{(z-\mu)^2}{2\sigma^2})$ gives the frequency of each phenotype $z$ assuming a normally distributed population. Thus, the product $w(z) \bullet f(z)$ weights the fitness of each phenotype ( $w(z)$) by its expected frequency within the population ($f(z)$) to calculate the expected mean survival across the population. The integral was bounded between $\theta - 100$ and $\theta + 100$ because the numerical integration algorithm used in R failed with infinite bounds when θ was far from 0; these bounds approximated the infinite integral while ensuring we captured the relevant variation in the functions.
